# Retromer Combinatorials for Gene-Therapy Across a Spectrum of Neurological Diseases

**DOI:** 10.1101/2020.09.03.282327

**Authors:** Yasir H. Qureshi, Vivek M. Patel, Suvarnambiga Kannan, Samuel D Waksal, Gregory A. Petsko, Scott A. Small

## Abstract

Endosomal trafficking is a biological pathway implicated in Alzheimer’s and Parkinson’s disease, and a growing number of other neurological disorders. For this category of diseases, the endosome’s trafficking complex retromer has emerged as a validated therapeutic target. Retromer’s core is a heterotrimeric complex composed of the scaffold protein VPS35 to which VPS26 and VPS29 bind. Unless it is deficient, increasing expression of VPS35 by viral vectors has a limited effect on other trimeric members and on retromer’s overall function. Here we set out to address these constraints and, based on prior insight, hypothesized that co-expressing VPS35 and VPS26 would synergistically interact and elevate retromer’s trimeric expression and function. Neurons, however, are distinct in expressing two VPS26 paralogs, VPS26a and VPS26b, and so to test the hypothesis we generated three novel AAV9 vectors harboring the VPS35, or VPS26a, or VPS26b transgene. First, we optimized their expression in neuroblastoma cell lines, then, in a comprehensive series of neuronal culture experiments, we expressed VPS35, VPS26a, and VPS26b individually and in all possible combinations. Confirming our hypothesis, expressing individual proteins failed to affect the trimer, while VPS35 and VPS26 combinatorials synergized the trimer’s expression. In addition, we illustrate functional synergy by showing that only VPS35 and VPS26 combinatorials significantly increase levels of Sorl1, a key retromer-receptor deficient in Alzheimer’s disease. Collectively, and together with other recent observations, these results suggest a precision-medicine logic when applying retromer gene therapy to a host of neurological disorders, depending on each disorder’s specific retromer-related molecular and anatomical phenotype.

## INTRODUCTION

Endosomal trafficking has emerged as a unified biological pathway disrupted in a growing number of neurological disorders. While clearly pathogenic in Alzheimer’s disease (AD) [1, 2] and Parkinson’s disease (PD) [3], endosomal trafficking defects are now thought to be central to disorders as varied as neuronal ceroid lipofuscinosis (NCL) [4, 5], Down’s syndrome [6], hereditary spastic paraplegia (HSP) [7], prion disease [8], and amyotrophic lateral sclerosis (ALS)-frontal-temporal degeneration (FTD) [9].

Retromer is a protein assembly that is a ‘master conductor’ of endosomal trafficking [10, 11], which functions as a molecular machine dedicated to recycling cargo out of endosomes. Perhaps unsurprisingly, nearly all neurological disorders in which endosomal trafficking has been implicated have links to retromer and its trafficking pathway [12-17]. Retromer is therefore considered a validated therapeutic target for neurological disorders, even the most common ones like Alzheimer’s and Parkinson’s [18].

The retromer assembly is organized into molecularly and functionally distinct modules that work in unison. Despite retromer’s complexity, previous studies have identified how it can be therapeutically targeted. Retromer’s ‘cargo recognition’ core is considered the machine’s central regulator, since it is with this core that all other retromer modules interact [18]. Thus, a pharmacological assumption is that targeting retromer’s core might be sufficient to enhance the function of the whole molecular machine, thereby increasing endosomal trafficking. This assumption and its therapeutic utility were first validated with the identification of ‘retromer pharmacological chaperones’ [19]. The retromer core is a heterotrimeric complex, with ‘vacuolar protein sorting 35’ (VPS35) at its center, to which VPS29 and VPS26 bind at either of its ends (Fig 1A). Disrupting the binding of VPS35 with other retromer core proteins leads to their accelerated degradation *in vitro* [19] and *in vivo* [20], justifying a search for retromer chaperones that would strengthen this interaction. Two such small molecule chaperones were isolated in a series of screens, and when applied to neuronal culture, the chaperones were found to increase the levels of multiple retromer core proteins [19]. Validating that an increase in retromer core proteins is associated with an increase in retromer’s overall function, the chaperones were also found to enhance the recycling of one of the brain’s key retromer-receptors, Sorl1 [19]. Because loss-of-function mutations in *SORL1* are causal in Alzheimer’s disease [21], and an approximate 30% reduction in Sorl1 protein is found even in early stages of the sporadic disease [22-24], the relatively modest effect on Sorl1 recycling was deemed therapeutically meaningful. Since then multiple independent studies have shown that the chaperones meaningfully benefit retromer’s trafficking function[25-29], including in human neurons, flies and mice, which collectively validate the concept that increasing retromer core protein levels is sufficient to functionally enhance endosomal trafficking.

**Figure 1.**
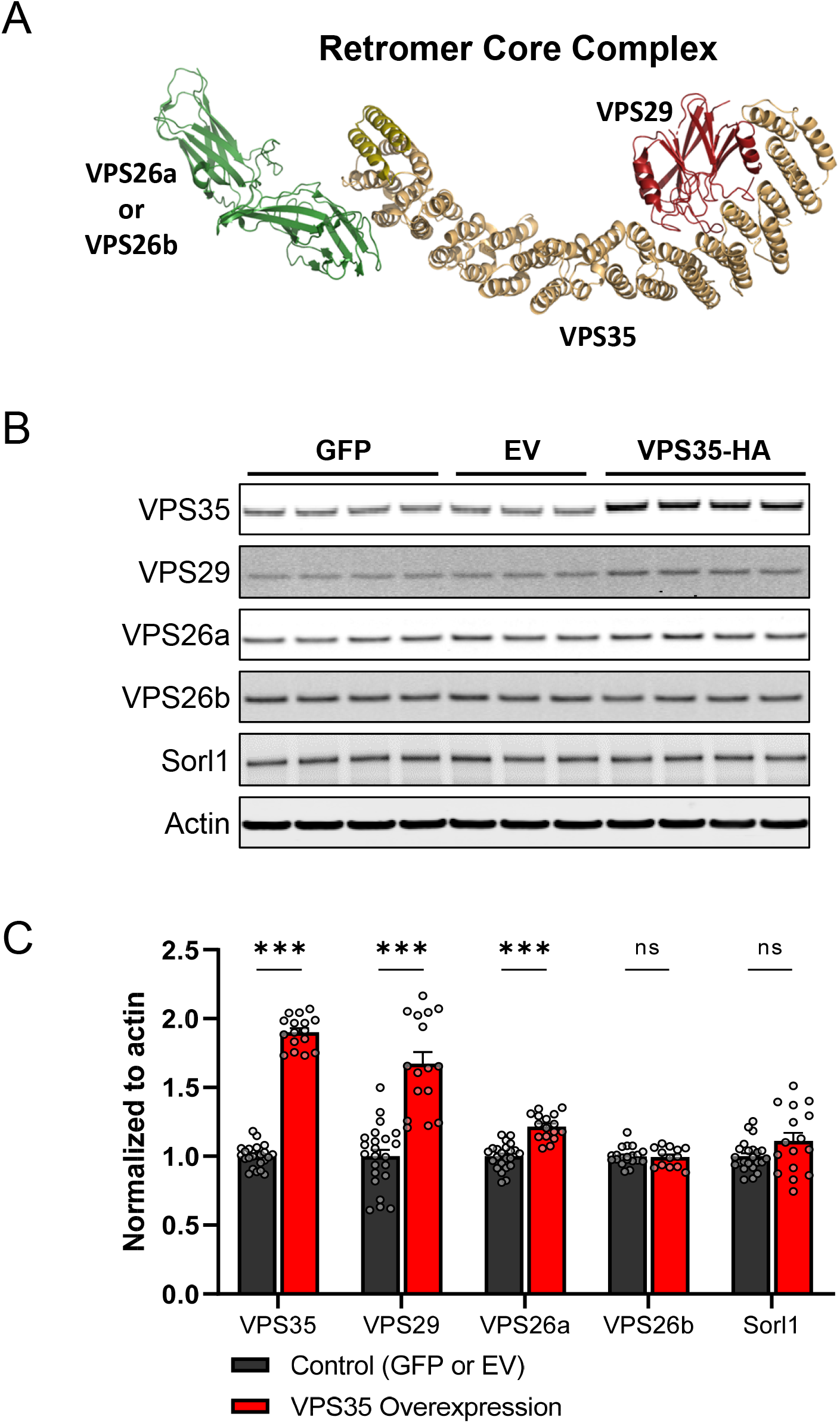
VPS35 expression alone is insufficient to elevate retromer’s trimer and function. (A) Backbone rendering (PyMOL, Schroedinger, Inc.) of the three-dimensional structure [55] of retromer’s cargo recognition core highlighting the interaction of VPS35 (orange) with VPS29 (red) and VPS26a (green); Vps26b binds to Vps35 in a virtually identical fashion [56, 57]. Atomic coordinates (PDB file 6vac) taken from the cryoEM structure of the mouse heterotrimer [55]. (B) Representative immunoblots showing retromer and Sorl1 expression levels following AAV9-VPS35-HA. AAV9-GFP and AAV9-EV (empty vector) were used as controls. (C) Bar graphs show mean levels of retromer components and Sorl1 from B, normalized to actin; Control (dark grey, n=23), VPS35 Overexpression (red, n=16). ***P < 0.001, ns= not significant.

If increasing retromer core protein levels is beneficial, then an inescapable conclusion is that, besides pharmacological chaperones, gene therapy might expand the retromer pharmacopeia. Indeed, we [30] and others [31] have used viral vector technology to show that expressing exogenous VPS35 in mouse models of AD can ameliorate disease-associated pathology. These mouse models, however, are VPS35 deficient [27, 30]. Many neurological disorders would benefit from enhancing retromer-dependent endosomal trafficking, even if VPS35 is not genetically altered or deficient.

For example, the approximately 30% Sorl1 deficiency observed in Alzheimer’s disease brains may occur independent of retromer levels.

While VPS35 is central to the core, the effects of overexpressing it alone will be limited by the availability of its other core partners. Based on recent ribosome profiling and proteomic studies [32-34], the levels of VPS29 are thought to be roughly double of those of VPS26 in virtually all cells, implying that VPS26 might be a limiting component. Additionally, an independent series of *in vitro* and cell biological observations suggest that VPS26 might interact with VPS35 in further stabilizing the trimer [19, 35]. If so, a synergistic effect of VPS26 expression might be expected. In the context of neurological disorders, testing this idea is complicated by the fact that the brain is distinct in expressing two VPS26 paralogs. Differentiating the two, the first, ubiquitously expressed paralog is called VPS26a, while the brain enriched paralog is called VPS26b [36]. Thus, to test our hypothesis we performed a study in cultured neurons in which all three (VPS35, VPS26a and VPS26b) are delivered by AAV9 and expressed individually and in all possible combinations: VPS35 alone, VPS26a alone, VPS26b alone; or, VPS35+VPS26a, VPS35+VPS26b, VPS26b+VPS26a, VPS35+VPS26a+VPS26b.

A detailed analysis from the comprehensive dataset validated the primary hypothesis, showing that only VPS35 and VPS26 combinatorials synergizes the expression of each, namely above and beyond what is observed when expressing VPS35 or VPS26 individually, and the expression of VPS29. While previous studies have established that this degree of trimer overexpression is therapeutically beneficial, we also provide evidence that only combinatorials have a synergistic effect on retromer’s function, by showing that they cause a significant increase in Sorl1 levels, sufficient to replete the Sorl1 deficiency observed in AD [22-24]. When taken together with previous observations, our findings suggest a precision-medicine approach for retromer gene therapy, depending on the molecular and anatomical phenotype of the range of endosomal trafficking disorders of the brain.

## RESULTS

### VPS35 expression alone is insufficient to elevate retromer’s trimer and function

To determine the effect exogenous VPS35 overexpression has on retromer core proteins and on retromer function in a non-deficient state, we transduced cultured wild-type mouse neurons with AAV9-VPS35-HA and used either AAV9-GFP or AAV9-empty vector (EV) as control conditions, and harvested them 3 weeks later.

The levels of all retromer core proteins were determined by immunoblotting (Fig. 1B). Compared to controls, a 90% VPS35-HA overexpression leads to a robust 67% increase in endogenous VPS29 (p=9E-09), a small 22% increase in VPS26a (p=2E-08), and no increase in VPS26b (p=0.62) (Fig. 1C).

VPS10-containing transmembrane proteins are established retromer cargo, and among these Sorl1 is the one most strongly linked to the neuronal retromer. By recycling transmembrane proteins out of the endosome, retromer keeps them away from the degradation pathway, which is why regulating retromer function is linked to Sorl1 levels. Accordingly, we immunoblotted for Sorl1. Compared to controls, overexpression of VPS35 alone had a slight (11%) and statistically unreliable (p=0.06) increase in Sorl1 (Fig. 1C).

By showing that VPS35 overexpression leads to a robust overexpression of VPS29, but either no or a modest increase in VPS26 paralogs, and no clear effect on retromer function, these results justify investigating the effect of VPS35 and VPS26 co-expression.

### Optimizing combined VPS35 and VPS26 expression in neuroblastoma cells

Before testing the effect of VPS35 and VPS26 combinatorials in neuronal culture, which requires viral vector delivery and a month-long study, we first turned to mouse neuroblastoma (N2A) cell lines in which our newly developed plasmids of VPS35, VPS26a, or VPS26b could be tested more rapidly via transfection. While not the main purpose of this study, we also took this opportunity to explore whether VPS35 and VPS26 co-expression interact in regulating retromer core protein levels.

N2A cells were transfected with plasmids expressing single proteins (VPS35, VPS26a, or VPS26b), or a combination of proteins (VPS35+VPS26a or VPS35+VPS26b). A plasmid expressing GFP or an empty plasmid were used as controls (Fig. 2A). Single protein conditions resulted in overexpression of each protein above control levels: for VPS35 alone (80%, p=3.4E-09), for VPS26a alone (550%; p=2.2E-06), and for VPS26b alone (362%; p= 0.0002) (Fig. 2B). When compared to single protein conditions, VPS35+VPS26a expression resulted in a significant increase in VPS35 (31%; p=0.0003), VPS29 (17%; p=0.0007), and VPS26a (52%; p=0.015), but minimal change in VPS26b; and VPS35+VPS26b expression resulted in a non-significant increase in VPS35 (15%; p=0.07), and VPS26b (56%; p=0.14), a significant increase in VPS29 (22%; p=0.0005), but no increase in VPS26a (Fig. 2B).

**Figure 2.**
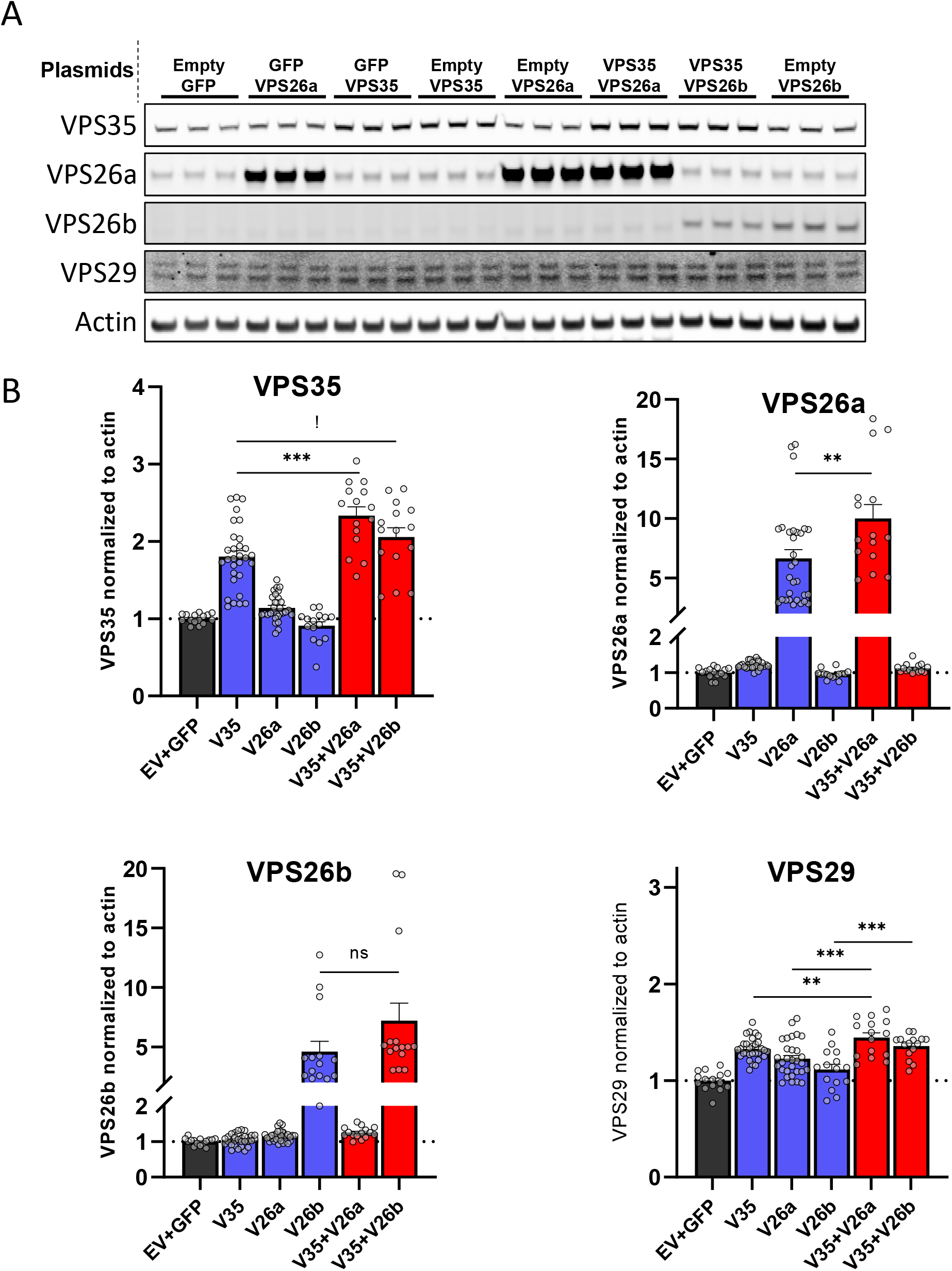
Optimizing combined VPS35 and VPS26 expression in neuroblastoma cells. (A) Representative immunoblots showing retromer expression levels following transfection with plasmids containing VPS35, VPS26a, and VPS26b alone or dual transfection of VPS35 along with VPS26a or VPS26b. GFP or empty backbone plasmids were used as controls. To properly control for the amount of plasmid DNA/lipofectamine complexes introduced in each condition, a control plasmid (GFP or empty backbone) was included whenever only one component of retromer was transfected. VPS29 shows 2 distinct bands, which represent two different isoforms of this protein in these N2a cells. (B) Bar graphs show mean levels of retromer components from A, normalized to actin; Control (dark grey, n=15), VPS35 (n=30), VPS26a (n=30), and VPS26b (n=15) overexpression alone (blue), VPS35 + VPS26 combinatorials (red, n=15). ***P < 0.001, **P <0.01, !P=0.07, ns= not significant.

### Combined VPS35 and VPS26 expression synergizes the retromer trimer in neurons

To test the effect of VPS35 and VPS26 combinatorials in cultured neurons, we generated five experimental AAV9 vectors, expressing mouse VPS35, VPS26a, VPS26b, and two control AAV9 vectors, one expressing GFP and the other an empty vector. The experimental vectors were expressed in neurons in all possible combinations: Single protein expression (VPS35 alone, VPS26a alone, VPS26b alone); double protein expression (VPS35+VPS26a, VPS35+VPS26b, VPS26b+VPS26a); and triple protein expression (VPS35+VPS26a+VPS26b) (Fig. 3A).

**Figure 3.**
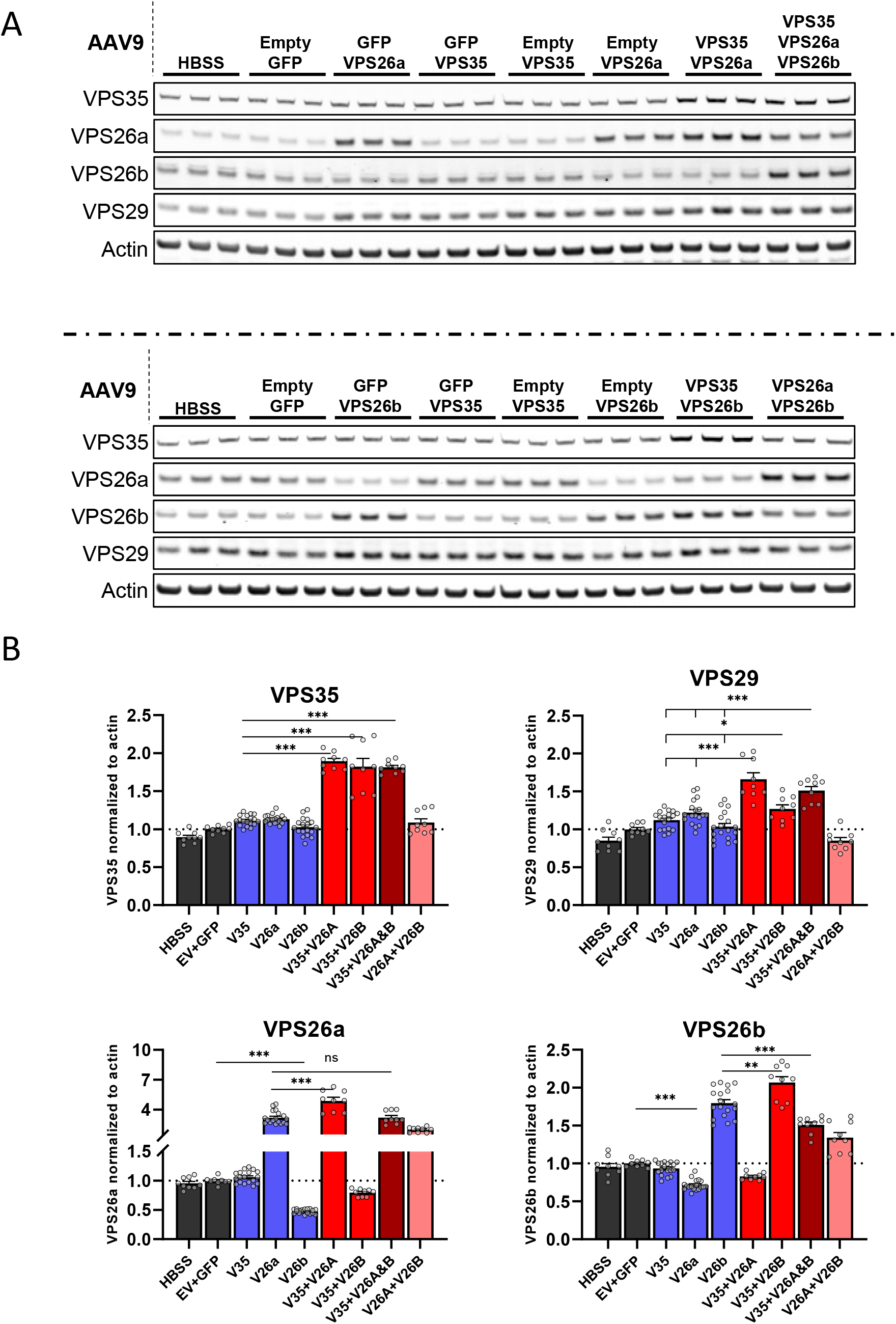
Combined VPS35 and VPS26 expression synergizes retromer trimer in neurons. (A) Representative immunoblots showing retromer expression levels following transduction with AAV9 vectors containing VPS35, VPS26a, and VPS26b. AAV9-GFP and AAV9-EV (AAV9 containing empty backbone plasmid) were used as controls. The experimental AAV9 vectors were expressed in neurons in all possible combinations: Single protein expression (VPS35 alone, VPS26a alone, VPS26b alone); double protein expression (VPS35+VPS26a, VPS35+VPS26b, VPS26b+VPS26a); and triple protein expression (VPS35+VPS26a+VPS26b). To properly control for the amount of DNA & AAV9 introduced in each condition, a control AAV9 (GFP or EV) was included whenever only one component of retromer was transduced. (B) Bar graphs show mean levels of retromer components from A, normalized to actin; Control (dark grey, n=9), single overexpression of VPS35, VPS26a, or VPS26b (blue, n=18), VPS35 + VPS26a or VPS26b combinatorials (red, n=9), VPS35 + VPS26a + VPS26b (dark red, n=9), VPS26a + VPS26b (pink, n=9). ***P < 0.001, **P < 0.01, *P < 0.05, ns= not significant.

The dose of each viral vector was optimized in exploratory studies, and when used in the final combinatorial study, mean AAV9-VPS35 overexpression was 11% (range:-1% to 23%), mean AAV9-VPS26a was 218% (range: 154% to 354%) and mean AAV9-VPS26b was 80% (range: 50% to 107%) (Fig. 3B, blue bars). This profile turned out to be particularly useful for testing for synergistic interactions.

We first tested whether there was a VPS35-VPS26 interaction on retromer core protein expression, by comparing levels detected in single protein conditions to those detected in the combinatorial experiments. Compared to single protein expression, VPS35+VPS26a expression resulted in a significant increase in VPS35 (70%; p=4.4E-18), VPS26a (53%; p=2.1E-05), VPS29 (∼42%; p<1.22E-05), but not VPS26b. VPS35+VPS26b expression resulted in a significant increase in VPS35 (64%; p=3.6E-09), VPS26b (15%; p=0.003), VPS29 (∼18%; p<0.013), but not VPS26a. Finally, VPS35+VPS26a+VPS26b expression resulted in a significant increase in all four retromer proteins compared to controls (EV+GFP) -- VPS35 (81%; p=5.6E-15), VPS29 (51%; p<1.8E-07), VPS26a (220%; p=1.7E-08) and VPS26b (51%; p=9.8E-10) (Fig. 3B).

While the main purpose of this comprehensive series of experiments was to test for synergistic interactions, the fact that VPS35+VPS26a did not increase VPS26b, and that VPS35+VPS26b did not increase VPS26a, suggests that in neurons each VPS26 paralog exists in biochemically distinct trimers.

### Combined VPS35 and VPS26 expression synergizes retromer function in neurons

We then tested whether VPS35 and VPS26 combinatorials have a synergistic effect on retromer function, by comparing levels of Sorl1 measured across all conditions. We used a univariate ANOVA, in which control conditions, single conditions, and combinatorial conditions were included as the fixed factor, and Sorl1 was included as the dependent variable. Results revealed a group effect (F= 19.3, p= 9.3E-8), with a simple comparison indicating that, while there was no difference between the control and single conditions (contrast estimate=0.1, p= 0.9), there was a significant difference between the single and combination conditions (contrast estimate=0.4, p=2.2 E-7). Post-hoc comparisons revealed that each combination condition was significantly different than the single conditions (Supplementary Fig. 1).

Interestingly, we also observed a significant effect of VPS26a+VPS26b overexpression on Sorl1 (34%; p=0.006) (Supplementary Fig.1), even though this combination condition did not increase levels of VPS35 or VPS29 (Fig. 3B). Sorl1 has been found to interact with the retromer core via VPS26 [37] (Fig. 4c), and our observation suggests that both paralogs might independently interact with Sorl1, which is also implied by their structures (Fig.4c).

**Figure 4.**
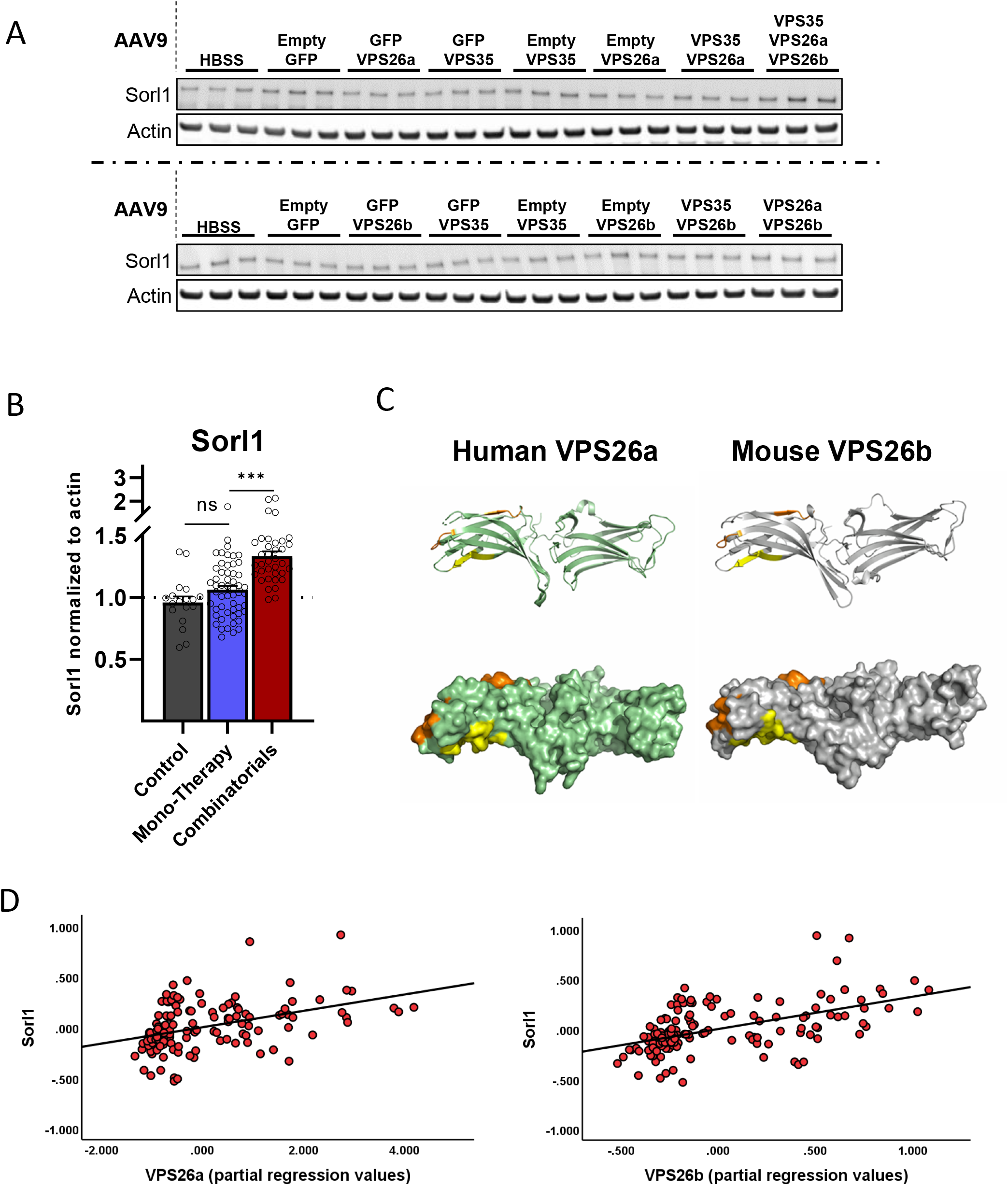
Combined VPS35 and VPS26 expression synergizes retromer function in neurons. (A) Representative immunoblots showing Sorl1 expression levels following transduction with AAV9 vectors containing VPS35, VPS26a, and VPS26b. AAV9-GFP and AAV9-EV (AAV9 containing empty backbone plasmid) were used as controls. The experimental AAV9 vectors were expressed in neurons in all possible combinations (details in Fig. 3). (B) Bar graph show mean levels of Sorl1 from A, normalized to actin; Control (dark grey, n=18), single overexpression of VPS35, VPS26a, or VPS26b (blue, n=54), combinatorials (dark red, n=36) ***P < 0.001, ns= not significant. (C) Ribbon and space-filing rendering (PyMOL, Schroedinger, Inc.) of the three-dimensional structures of retromer component paralogs VPS26a (left, PDB coordinates 2FAU, ref. [57]) and VPS26b (right, PDVB coordinates 2R51, ref. [56]), with highlighted orange and yellow residues that have been implicated in cargo recognition in various studies. The orange residues have been identified [37] as being essential for binding of the yeast Sorl1 orthologue VPS10. These residues are almost completely conserved from yeast to humans [37]. (D) Scatter plots generated from a multivariate regression, demonstrating that VPS26a (t=5.6, p=1.4E-7) and VPS26b (F=7.2, p=3.2E-11) independently correlate with Sorl1 levels.

We took advantage of the large-scale dataset we generated, in which there are over 140 experimental or control conditions and in which the four retromer core proteins and Sorl1 were measured across a broad dynamic range. A multiple linear regression model was used, in which Sorl1 levels was included as the dependent variable, and VPS35, VPS26, VPS26a and VPS26b levels were simultaneously entered as the independent variables. A significant relationship to Sorl1 levels was found (F=19.5, p=1.1E-12), with only the VPS26 paralogs significantly contributing to the model. The model was therefore trimmed to include both paralogs, confirming that both VPS26a (t=5.6, p=1.4E-7) and VPS26b (F=7.2, p=3.2E-11) independently correlate with Sorl1 levels (Fig. 4B). This result agrees with the interpretation that neurons have two trimers (VPS26b-VPS35-VPS29 and VPS26a-VPS35-VPS29) that are not only biochemically distinct, but functionally distinct as well.

## DISCUSSION

Previous studies in non-neuronal cells have suggested that there is a stoichiometric imbalance among the three retromer core proteins, with VPS29 level being nearly double that of VPS26 [32-34]. Our first study, in which we overexpressed exogenous VPS35, provides independent validation, and suggests that the same VPS29-VPS26 mismatch exists in neurons. In our previous work with AAV9-VPS35, we found that VPS35 is tightly autoregulated [30], a result also obtained in HeLa cells in culture [20], so that with increased exogenous VPS35 expression there is an accompanying and comparable decrease in endogenous VPS35. This autoregulation is due to the relative instability of VPS35 in the cell when it is not bound to the other proteins of the heterotrimer [20] (it is also the least stable of the core proteins to thermal denaturation *in vitro* [19, 35]).

When overexpression is ultimately achieved, nearly all VPS35 in the cell is replaced by its exogenous form, and we have found that this exogenous VPS35 binds its endogenous core partners and forms a functional retromer heterotrimer [30]. These observations help interpret our first study, in which overexpressed exogenous VPS35 can be effectively used to probe for the levels of its endogenous partners. Finding that VPS35 overexpression had a much more dramatic effect in increasing levels of endogenous VPS29 than endogenous VPS26, supports previous inference on the trimer’s stoichiometry and assembly, and is broadly consistent with the VPS29-VPS35 heterodimer being a stable assembly intermediate [35]. Finding that VPS35 and VPS29 overexpression had no effect on retromer function, relying on an established cargo of the neuronal retromer, confirms the conclusion from previous studies, that all three core proteins need to be elevated to upregulate retromer function, a conclusion also supported by the three-dimensional structure of the retromer core complex [38].

While informative, the purpose of this study was not to show that when overexpressing VPS35, VPS26 is retromer’s limiting component. The main motivation was to test the hypothesis that VPS35 and VPS26 actively interact *in vivo* in a way that would synergize to elevate all the components of the trimer. The hypothesis is based in part on *in vitro* studies, in which we showed that, when they bind one another, VPS26 stabilizes the longer and much less stable VPS35 [19]. Retromer’s structural biology might also account for this synergism, with VPS26 effectively acting as the stable docking site for binding of VPS35 and the VPS35-VPS29 heterodimer [38]. Testing this hypothesis in neurons is complicated by the fact that the brain is distinct in expressing two VPS26 paralogs—VPS26a and VPS26b [36]. Thus, in order to formally test for an interaction, multiple combinatorials needed to be evaluated. We started in N2A cells to optimize the combinatorials, and even in this cell line we observed evidence in support of an interaction. The stronger evidence came from the ultimate study in neurons, in which we started with low exogenous VPS35 expression, and used all 7 combinatorials of VPS35, VPS26a, and VPS26b. By comparing the expression profiles of VPS35 and the VPS26 paralogs, when each were expressed alone or in combination, we provide clear evidence of a non-additive interaction-- which is to say synergism-- in elevating all components of the trimer.

Evidence for an interaction is also implied by our readout of retromer function, an increase in Sorl1 levels by approximately 34%, an effect size that mirrors the degree of Sorl1 deficiency observed in AD [22-24]. A broader comparison on the effect of Sorl1 across the studies informs on retromer functionality. In the first neuronal study, when VPS35 was overexpressed alone, two out of the three retromer core proteins were robustly elevated, but with no effect on retromer function. In the second neuronal study, when thanks to synergism all three of the trimeric proteins were robustly elevated, this led to an increase in retromer function. This result supports the conclusion that all three retromer core proteins need to be co-elevated to upregulate retromer’s overall function. In fact, this conclusion is not at all obvious. From protein abundance considerations, since VPS29 is always considerably in excess under normal cellular conditions, one would naively expect that only VPS26 and VPS35 need to be elevated. Our studies show that by exploiting retromer’s stoichiometry and protein-protein interactions there is no need to exogenously express all three proteins. Exogenously expressing VPS35 and VPS26 is sufficient to also upregulate the level of VPS29 and increase retromer’s endosomal cargo recycling function. We also show, importantly, that the levels of VPS26a and VPS26b are independent of one another, and therefore appropriate choice of combinatorials allows selective increase of one retromer heterotrimer over the other.

Collectively, together with other recent studies [30], these findings suggest a precision-medicine approach for using gene therapy across a host of neurological disorders, depending on their retromer-related molecular and anatomical phenotypes (Table). In rare cases in which deleterious mutations in VPS35 are identified, as they sometimes are in PD [39] and rarely in AD [40], viral vectors expressing ‘wildtype’ exogenous VPS35 alone might be ideal. The fact that endogenous VPS35 is tightly autoregulated can be exploited because high enough exogenous wild type VPS35 expression will replace the endogenous protein, effectively acting as VPS35 ‘replacement therapy’.

**Table.**
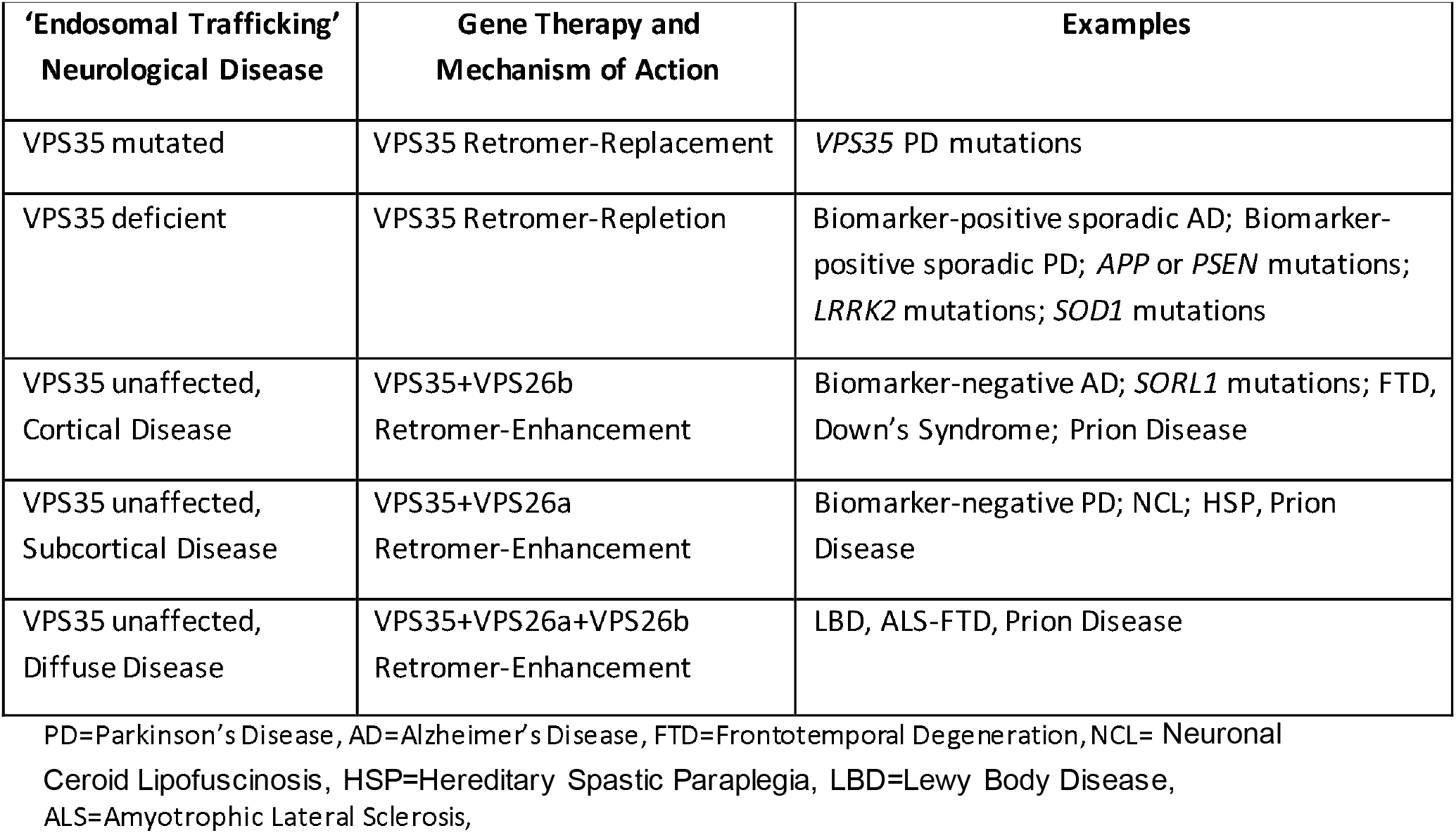
Retromer Gene Therapy and Precision Medicine

In cases where VPS35 is not mutated but the protein is nevertheless deficient, VPS35 monotherapy alone might still be sufficient, acting as ‘retromer-repletion’ therapy (Table). VPS35 deficiency can occur secondarily to monogenic disease-causing mutations, such as *LRRK2* mutations in PD [41], *APP* and *PSEN* mutations in AD [27, 42], and *SOD1* mutations in ALS [17]. Besides these monogenic forms of neurodegeneration, VPS35 deficiency has also been found in vulnerable regions of ‘sporadic’ forms of AD[12], PD[43] and ALS[17]. Because ‘sporadic’, for these disorders biomarkers are required to precisely determine which patients have evidence of VPS35 deficiency. A recent study, for example, identified a CSF-based biomarker of VPS35 deficiency in cortical neurons, and used this biomarker to identify a subset of sporadic AD patients with evidence of VPS35 deficiency [44]. In the case of sporadic PD, reduction in striatal dopamine transporter (DAT), as detected in vivo by SPECT and PET, is considered a reliable biomarker [45, 46]. Retromer turns out to be required for recycling striatal DAT[47], and VPS35 mutations and deficiencies have been shown to cause a reduction in striatal DAT [48]. While striatal neuronal death can act as the cause of DAT signal loss in late stages of PD, DAT reduction might be a biomarker of VPS35 deficiency in the disease’s earlier stages.

Enhancing retromer-dependent endosomal trafficking, however, is now thought to confer a therapeutic benefit across many neurological conditions even when VPS35 is not mutated or deficient, For example, since retromer enhancement has been shown to be beneficial in model systems of AD [26] and PD [25] even when VPS35 is unaffected, ‘retromer-enhancing’ gene therapy is expected to be beneficial even in those biomarker-negative AD and PD patients. Additionally, the list of diseases that might benefit from ‘retromer enhancement’ therapy includes FTD, NCL, HSP, Down’s syndrome, and some forms of prion disease.

For this broadest category of endosomal trafficking disorders, our results establish that co-expressing VPS35 with VPS26 is an efficient and reliable ‘retromer-enhancement’ gene therapy strategy (Table). In these cases, the complexity of the neuronal retromer, comprised of either VPS26a or VPS26b, can be utilized together with anatomical biology to allow the logic of precision-medicine to be applied. Gene expression atlases (Allen Human Brain Atlas. Available from: human.brain-map.org) suggest that VPS26b is differentially expressed in the cortex, while VPS26a is differentially expressed in the subcortex (Supplementary Fig. 2). Accordingly, VPS35+VPS26b retromer-enhancement would be most suitable for cortical endosomal trafficking disorders in which VPS35 is unaffected, with examples including biomarker-negative sporadic AD, AD patients with *SORL1* mutations, FTD, and Down’s syndrome. In contrast, VPS35+VPS26a retromer-enhancement is better suited for disorders in which endosomal trafficking defects occur primarily in subcortical regions, such as biomarker-negative sporadic PD, HSP, and some forms of NCL. Finally, VPS35+VPS26a+VPS26b might be indicated for endosomal trafficking neurological disorders in which the disease is more diffuse, such as Lewy Body Disease (LBD) and ALS-FTD.

## METHODS

### Design and preparation of the retromer constructs and AAV9

For initial experiments involving VPS35 expression, the DNA construct expressing the human VPS35 protein tagged with HA at the C-terminus was designed and synthesized using Thermofisher’s GeneArt portal. mRNA sequence for human VPS35 was acquired from the National Center for Biotechnology Information (NCBI). The construct was then subcloned into pcDNA 3.1 (+) Hygro vectors. For the control plasmids, EGFP sequence was obtained from pcDNA3-EGFP plasmid (Addgene plasmid 13031). DNA constructs were eventually subcloned into AAV9 transfer plasmids for viral production [49]. The expression cassette was flanked with adeno-associated virus serotype 2 (AAV2) terminal repeats. The cytomegalovirus chicken beta-actin hybrid promoter with CMV enhancer (“CAG” promoter) was used. The transgene was followed by the woodchuck hepatitis virus post-transcriptional regulatory element (WPRE) and the bovine growth hormone polyadenylation signal. Empty vector control was generated by deleting the GFP sequence. VPS35-HA transgene was subcloned into the construct in lieu of the GFP. Each of these constructs was individually packaged into a recombinant adeno-associated virus vector, AAV9, by described methods [50] using capsid and helper plasmid DNA from the University of Pennsylvania [51]. The AAV9 preparations were filter-sterilized using Millex syringe filters (Millipore) at the end of the procedure and then stored frozen in aliquots. The titering method was for encapsidated vector genomes per ml by a slot-blot method against a standard curve using the Amersham ECL Direct Nucleic Acid Labeling and Detection Systems (GE Healthcare Bio-Sciences).

For further experiments involving VPS26a, VPS26b, and VPS35 combinatorials, new plasmids were designed in collaboration with MeiraGTx for production of AAV vectors. Mouse mRNA sequences were acquired from the National Center for Biotechnology Information (NCBI). The sequences were codon optimized and then *de novo* synthesized. The synthesized construct was sub-cloned into AAV transfer plasmid with AAV2 inverted terminal repeats (ITRs) and a ubiquitous Chicken beta Actin wild type promoter. The transgene was followed by the bovine growth hormone polyadenylation signal. Empty chassis vector control was generated by deleting the VPS35 sequence. The GFP control was designed to mimic the rationale design of the target VPS-constructs. It contains an enhanced green fluorescent protein, plus the same Bovine Growth Hormone polyA (BgH) and Chicken beta Actin wild type promoter, as the VPS constructs. These plasmids were used to generate AAV9 retromer vectors at MeiraGTx labs. Each of the constructs described above were individually packaged into a recombinant adeno-associated virus vector 9 (AAV9), using capsid and helper plasmid DNA from MeiraGTx. Briefly, the transfer plasmid, rep-cap plasmid and helper plasmids were co-transfected into HEK 293 cells. The harvested suspension containing virus and cellular debris was clarified using millipore SHC XL150 filter (140cm2). The clarified suspension was then purified with AVB Sepharose, 20mL column, elution in 3 column volumes. Concentration and diafiltration was performed with 100kD mPES hollow fibre (Spectrum MicroKros cat# C02-E100-05-S). Further concentration was done with Amicon Ultra-4 Centrifugal Filter 30kD (cat# UFC8030).

### N2A Culture

Mouse neuroblastoma (N2a) cells were cultured in 50% DMEM (high glucose) & 50% Opti-MEM + 10% FBS and Glutamine (2mM) with penicillin and streptomycin to prevent microbial contamination.

### Transfection

Lipofectamine transfection protocols were used with some modifications. Briefly we used lipofectamine LTX to co-transfect VPS35 and VPS26 (VPS26a or VPS26b) plasmids into neuron like cells, Neuro2a (N2a) in a 6 well format. 100k cells were plated in each well already containing medium with DNA-lipofectamine complexes. We used empty chassis and GFP as control plasmids. Amount of DNA copies introduced per well was 2.81E+11. Cells were harvested 48 hours after transfection using RIPA buffer as described previously[30].

### Neuronal culture and transduction

Primary mouse cortical and hippocampal neuronal cultures were implemented as described previously [52]. Neurons were transduced with retromer AAV9 (Multiplicity of infection (MOI) of 83k, 67k, or 50k depending on retromer construct), 7 days after plating. Empty chassis AAV9 and GFP AAV9 were used as controls. The culture was maintained for 3 weeks after transduction (4 weeks total). At day 28 the neurons were lysed using RIPA buffer with protease and phosphatase inhibitors.

### Western blots

Cells from N2A and neuronal cultures were lysed in RIPA and proteins were isolated as described previously [30, 53]. Lysates from the samples were run on NuPAGE® Bis-Tris 4-12% gels, transferred onto nitrocellulose membranes using iblot and were probed with antibodies. Primary antibodies targeting the following proteins were used; VPS35 (ab57632, Abcam, 1:1k), VPS26a (ab211530, Abcam, 1:500), VPS26b (NBP1-92575, Novus, 1:500 or 15915-1-AP, proteintech, 1:500), VPS29 (sab2501105, Sigma-Aldrich, 1:500), Sorl1 (611861, BD-biosciences, 1:2k & 79322, Cell Signaling, 1:500), and β-actin (ab6276, Abcam, 1:5k). IRDye® 800 or 680 antibodies (LI-COR) were used as secondary with dilutions of 1:10k for 800CW, 1:15k for 680RD, and 1:25k for 680LT antibodies. Western blots were scanned using the Odyssey imaging system as described previously [54]. For Sorl1 (BD-611861) Peroxidase AffiniPure Donkey Anti-Mouse IgG (H+L) was used as secondary antibody (Jackson Immuno Research labs, 1:2k), and blots scanned at Fujifilm LAS-3000 Imager.

### Statistics

Statistical analysis was performed using Microsoft Excel and SPSS. Independent two-sample student’s t-test, assuming equal variance, with two-tailed distribution was used for all experiments unless stated otherwise. All data are presented as means, the error bars indicate standard error of the mean. All bar graphs were created in GraphPad Prism 8. Scatter plots were created in SPSS.

## Acknowledgements

This study was partly supported by an NIH grant AG008702 and an anonymous foundation to S.A.S. and sponsored research from Meira GTX to G.A.P.

## Disclosures

S.A.S and G.A.P are on the Scientific Advisory Board of Meira GTX

### FIGURE LEGENDS

**Supplementary Figure 1.**
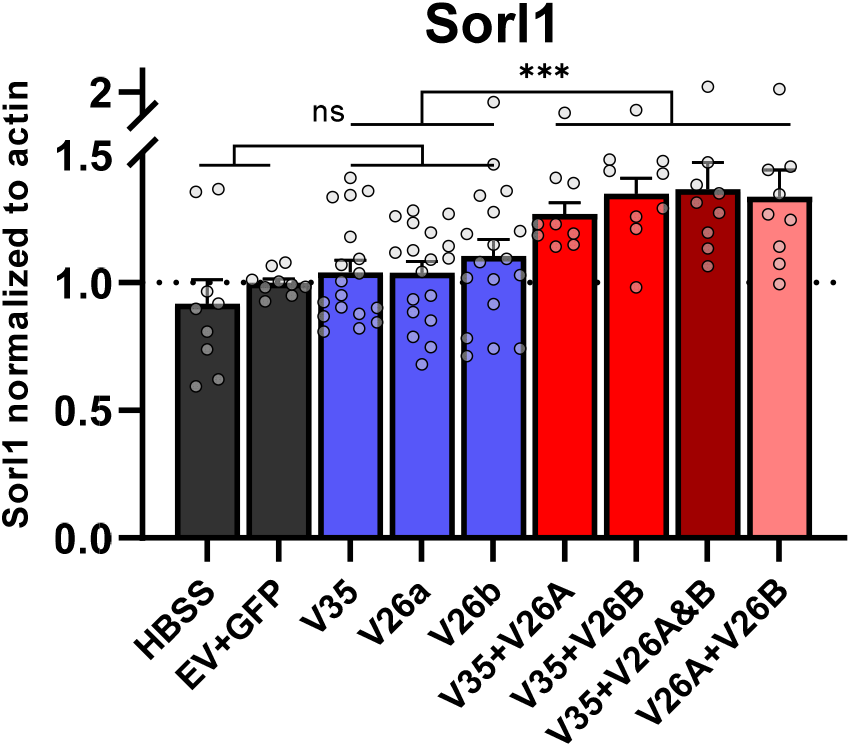
VPS35 and VPS26 combinatorials and Sorl1 expression. Bar graphs show mean levels of Sorl1 from Fig 4A, normalized to actin; Control (dark grey, n=9), single overexpression of VPS35, VPS26a, or VPS26b (blue, n=18), VPS35 + VPS26a or VPS26b combinatorials (red, n=9), VPS35 + VPS26a + VPS26b (dark red, n=9), VPS26a + VPS26b (pink, n=9). ***P < 0.001, ns= not significant.

**Supplementary Figure 2.**
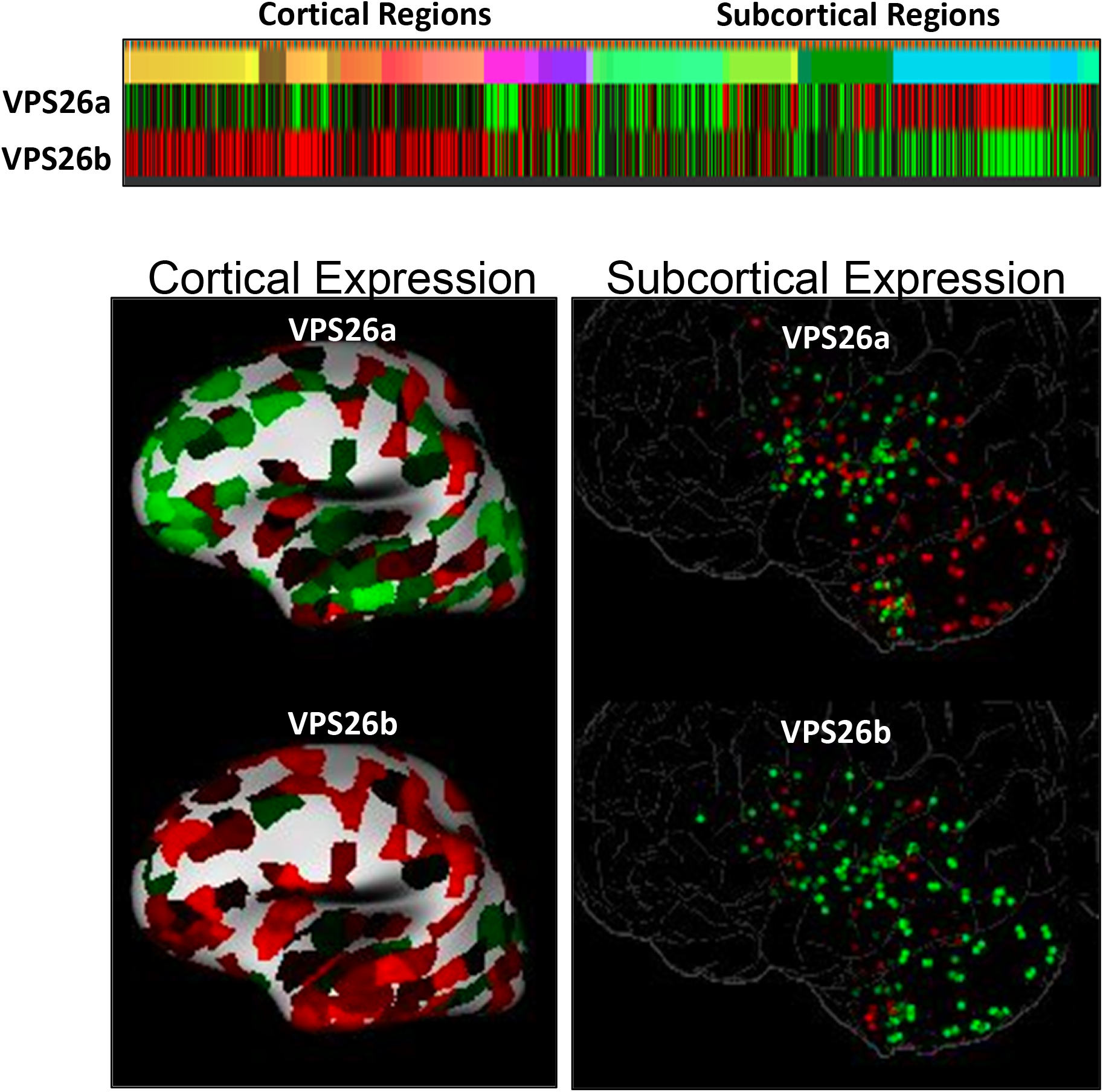
The anatomical biology VPS26. As documented in the Allen Human Brain Atlas (© 2010 Allen Institute for Brain Science. Allen Human Brain Atlas. Available from: human.brain-map.org), VPS26a is differentially expressed in subcortical regions, while VPS26b is differentially expressed in cortical regions. (Higher expression is indicated in red to lower expression in green).

